# Genomic and phenotypic analysis of COVID-19-associated pulmonary aspergillosis isolates of *Aspergillus fumigatus*

**DOI:** 10.1101/2020.11.06.371971

**Authors:** Jacob L. Steenwyk, Matthew E. Mead, Patrícia Alves de Castro, Clara Valero, André Damasio, Renato A. C. dos Santos, Abigail L. Labella, Yuanning Li, Sonja L. Knowles, Huzefa A. Raja, Nicholas H. Oberlies, Xiaofan Zhou, Oliver A. Cornely, Frieder Fuchs, Philipp Koehler, Gustavo H. Goldman, Antonis Rokas

**Affiliations:** Department of Biological Sciences, Vanderbilt University, Nashville, Tennessee, USA; Faculdade de Ciências Farmacêuticas de Ribeirão Preto, Universidade de São Paulo, Ribeirão Preto, Brazil; Institute of Biology, University of Campinas (UNICAMP), Campinas-SP, Brazil; Experimental Medicine Research Cluster (EMRC), University of Campinas (UNICAMP), Campinas-SP, Brazil; Department of Chemistry and Biochemistry, University of North Carolina at Greensboro, North Carolina 27402; Guangdong Laboratory for Lingnan Modern Agriculture, Guangdong Province Key Laboratory of Microbial Signals and Disease Control, Integrative Microbiology Research Centre, South China Agricultural University, Guangzhou, China; University of Cologne, Medical Faculty and University Hospital Cologne, Department I of Internal Medicine, Excellence Center for Medical Mycology (ECMM), Cologne, Germany; University of Cologne, Cologne Excellence Cluster on Cellular Stress Responses in Aging-Associated Diseases (CECAD), Cologne, Germany; ZKS Köln, Clinical Trials Centre Cologne, Cologne, Germany; German Center for Infection Research (DZIF), Partner Site Bonn□Cologne, Medical Faculty and University Hospital Cologne, University of Cologne, Cologne, Germany; Faculty of Medicine, Institute for Medical Microbiology, Immunology and Hygiene, University of Cologne, Cologne, Germany

**Keywords:** pathogenicity, co-infection, secondary infection, virulence factors, superinfection, acute respiratory distress syndrome, *Aspergillus*

## Abstract

The ongoing global pandemic caused by the severe acute respiratory syndrome coronavirus 2 (SARS-CoV-2) is responsible for the coronavirus disease 2019 (COVID-19) first described from Wuhan, China. A subset of COVID-19 patients has been reported to have acquired secondary infections by microbial pathogens, such as fungal opportunistic pathogens from the genus *Aspergillus*. To gain insight into COVID-19 associated pulmonary aspergillosis (CAPA), we analyzed the genomes and characterized the phenotypic profiles of four CAPA isolates of *Aspergillus fumigatus* obtained from patients treated in the area of North Rhine-Westphalia, Germany. By examining the mutational spectrum of single nucleotide polymorphisms, insertion-deletion polymorphisms, and copy number variants among 206 genes known to modulate *A. fumigatus* virulence, we found that CAPA isolate genomes do not exhibit major differences from the genome of the Af293 reference strain. By examining virulence in an invertebrate moth model, growth in the presence of osmotic, cell wall, and oxidative stressors, and the minimum inhibitory concentration of antifungal drugs, we found that CAPA isolates were generally, but not always, similar to *A. fumigatus* reference strains Af293 and CEA17. Notably, CAPA isolate D had more putative loss of function mutations in genes known to increase virulence when deleted (e.g., in the *FLEA* gene, which encodes a lectin recognized by macrophages). Moreover, CAPA isolate D was significantly more virulent than the other three CAPA isolates and the *A. fumigatus* reference strains tested. These findings expand our understanding of the genomic and phenotypic characteristics of isolates that cause CAPA.

## Introduction

On March 11, 2020, the World Health Organization declared the ongoing pandemic caused by SARS-CoV-2, which causes COVID-19, a global emergency (Sohrabi et al., 2020). Similar to other viral infections, patients may be more susceptible to microbial secondary infections, which can complicate disease management strategies and result in adverse patient outcomes (Brüggemann et al., 2020; Cox et al., 2020). For example, approximately one quarter of patients infected with the H1N1 influenza virus during the 2009 pandemic were also infected with bacteria or fungi (MacIntyre et al., 2018; Zhou et al., 2020). Among studies examining patients with COVID-19, ∼17% of individuals also have bacterial infections (Langford et al., 2020) and one study found that ∼40% of patients with severe COVID-19 pneumonia were also infected with filamentous fungi from the genus *Aspergillus* (Nasir et al., 2020). Another study reported that ∼26% of patients with acute respiratory distress syndrome-associated COVID-19 were also infected with *Aspergillus fumigatus* and had high rates of mortality (Koehler et al., 2020). Despite the prevalence microbial infections and their association with adverse patient outcomes, these secondary infections are only beginning to be understood.

Invasive pulmonary aspergillosis is caused by tissue infiltration of Aspergilli after inhalation of asexual spores (Figure 1); more than 250,000 aspergillosis infections are estimated to occur annually and have high mortality rates (Bongomin et al., 2017). The major etiological agent of aspergillosis is *A. fumigatus* (Latgé and Chamilos, 2019), although a few other *Aspergillus* species are also known to cause aspergillosis (Bastos et al., 2020; dos Santos et al., 2020b; Rokas et al., 2020; Steenwyk et al., 2020c). Numerous factors are known to be associated with *A. fumigatus* pathogenicity, including its ability to grow at the human body temperature (37°C) and withstand oxidative stress (Kamei and Watanabe, 2005; Tekaia and Latgé, 2005; Shwab et al., 2007; Losada et al., 2009; Abad et al., 2010; Grahl et al., 2012; Yin et al., 2013; Wiemann et al., 2014; Knox et al., 2016; Kowalski et al., 2019; Raffa and Keller, 2019; Blachowicz et al., 2020). Disease management of *A. fumigatus* is further complicated by resistance to antifungal drugs among strains (Chamilos and Kontoyiannis, 2005; Howard and Arendrup, 2011; Chowdhary et al., 2014; Sewell et al., 2019) Additionally, *A. fumigatus* strains have been previously shown to exhibit strain heterogeneity with respect to virulence and pathogenicity-associated traits (Kowalski et al., 2016; Keller, 2017; Kowalski et al., 2019; Ries et al., 2019; dos Santos et al., 2020b; Steenwyk et al., 2020d). However, it remains unclear whether the genomic and pathogenicity-related phenotypic characteristics of CAPA isolates are similar or distinct from those of previously studied clinical strains of *A. fumigatus*.

**Figure 1.**
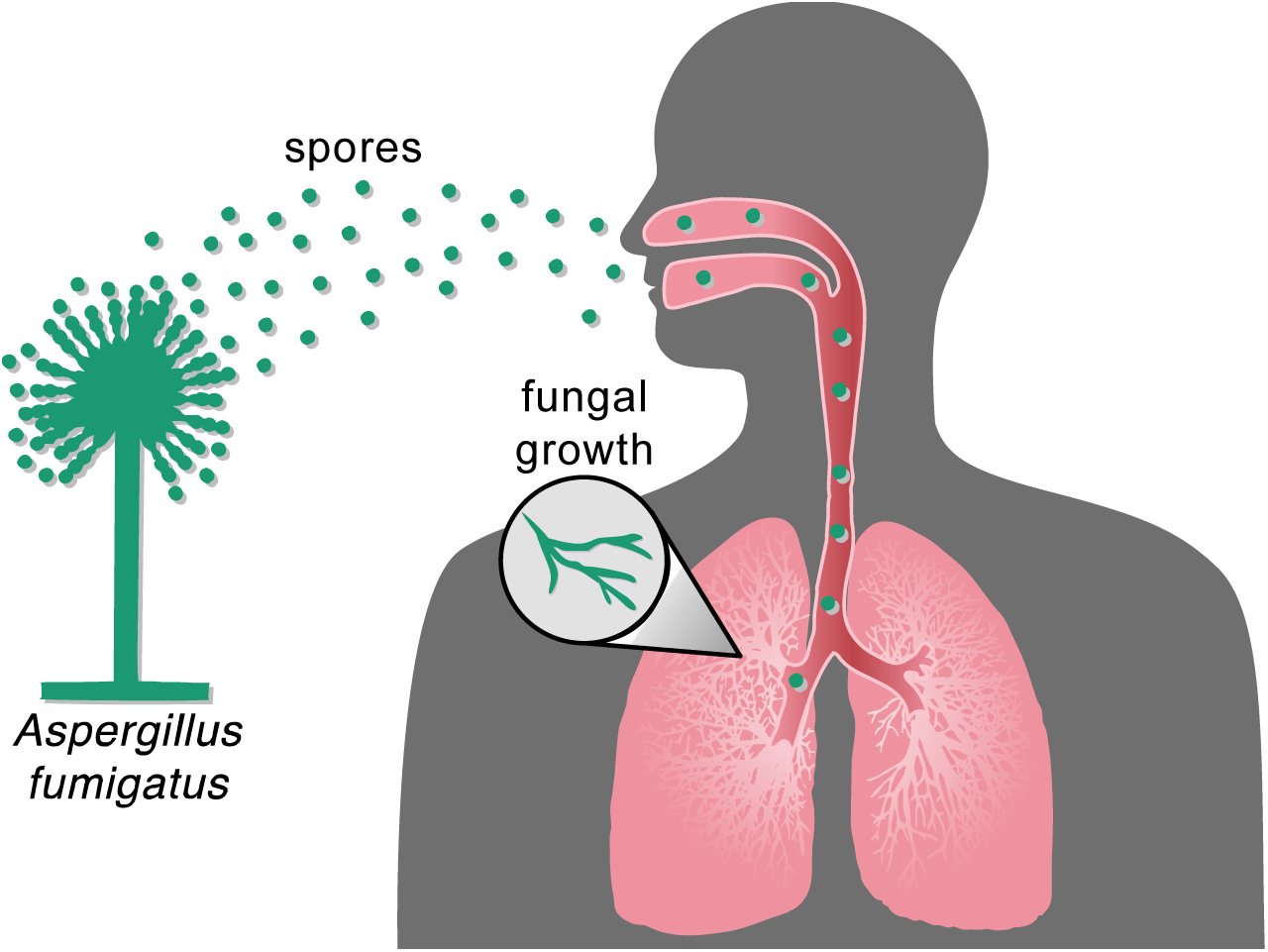
Inhalation of *Aspergillus* spores can result in fungal infection. Inhalation of *Aspergillus* spores from the environment can travel to the lung and then grow vegetatively and spread to other parts of the body.

To address this question and gain insight into the pathobiology of *A. fumigatus* CAPA isolates, we examined the genomic and phenotypic characteristics of four CAPA isolates obtained from four critically ill patients of two different centers in Cologne, Germany (Koehler et al., 2020) (Table 1). All patients were submitted to intensive care units due to moderate to severe respiratory distress syndrome (ARDS). Genome-scale phylogenetic (or phylogenomic) analyses revealed CAPA isolates formed a monophyletic group closely related to reference strains Af293 and A1163. Examination of the mutational spectra of 206 genes known to modulate virulence in *A. fumigatus* (which are hereafter referred to as genetic determinants of virulence) revealed several putative loss of function (LOF) mutations. Notably, CAPA isolate D had the most putative LOF mutations among genes whose null mutants are known to increase virulence. The profiles of pathogenicity-related traits of the CAPA isolates were similar to those of reference *A. fumigatus* strains Af293 and CEA17. One notable exception was that CAPA isolate D was significantly more virulent than other strains in an invertebrate model of disease. These results suggest that the genomes of *A. fumigatus* CAPA isolates contain nearly complete and intact repertoires of genetic determinants of virulence and have phenotypic profiles that are broadly expected for *A. fumigatus* clinical isolates. However, we did find evidence for genetic and phenotypic strain heterogeneity. These results suggest the CAPA isolates show similar virulence profiles as *A. fumigatus* clinical strains Af293 and A1163 and expand our understanding of CAPA.

**Table 1.** Metainformation and NCBI Accessions for CAPA isolates

| Isolate | Patient Identifier | Patient Outcome | Patient Age | Patient Sex | Patient immunocompromising condition | Antifungal treatment | Antiviral treatment | NCBI BioSample/Sequence Read Archive Accessions |
| --- | --- | --- | --- | --- | --- | --- | --- | --- |
| CAPA A | Patient from this study | Deceased | 57 | Male | None | Caspofungin (70/50 mg once daily) | Supportive only | SAMN16591136; SRR12949929 |
| CAPA B | Patient #4 | Deceased | 73 | Male | Inhalational steroids for medical history of chronic obstructive pulmonary disease | Voriconazole iv (6/4 mg/kg BW twice daily) | Supportive only | SAMN16591179; SRR12949928 |
| CAPA C | Patient #3 | Alive | 54 | Male | IV corticosteroid therapy 0.4 mg/kg/day, total of 13 days | Caspofungin (70/50 mg once daily) followed by IV voriconazole (6/4 mg/kg BW twice daily) | Hydroxychloroquine, darunavir and cobicistat at external hospital, in house changed to supportive only | SAMN16591190; SRR12949927 |
| CAPA D | Patient #1 | Deceased | 62 | Female | Inhalational steroids for medical history of chronic obstructive pulmonary disease | IV Voriconazole (6/4 mg/kg BW twice daily) | Supportive only | SAMN16591200; SRR12949926 |

Information on CAPA isolates B, C, and D is from (Koehler et al., 2020), where the isolates were first described. CAPA isolate A is being reported for the first time. Supportive only antiviral treatment indicates no specific antiviral treatment was given. Abbreviations are as follows: BW: body weight; IV: Intravenous; kg: kilogram; mg: milligram.

## Results and Discussion

### CAPA isolates belong to *A. fumigatus* and are closely related to reference strains Af293 and A1163

To confirm that the CAPA isolates belong to *A. fumigatus*, we sequenced, assembled, and annotated their genomes (Table S1). Phylogenetic analyses conducted using *tef1* (Figure S1) and calmodulin (Figure S2) sequences suggested that all CAPA isolates are *A. fumigatus*. Phylogenomic analysis using 50 *Aspergillus* genomes (the four CAPA isolates, 43 representative *A. fumigatus* genomes including strains Af293 and A1163 (Nierman et al., 2005; Fedorova et al., 2008; Liu et al., 2011; Abdolrasouli et al., 2015; Knox et al., 2016; Lind et al., 2017; Paul et al., 2017; dos Santos et al., 2020b), *A. fischeri* strains NRRL 181 and NRRL 4585, and *A. oerlinghausenensis* strain CBS 139183^T^ (Fedorova et al., 2008; Steenwyk et al., 2020d)) confirmed that all CAPA isolates are *A. fumigatus* (Figure 2A). Phylogenomic analyses also revealed the CAPA isolates formed a monophyletic group closely related to reference strains Af293 and A1163.

**Figure 2.**
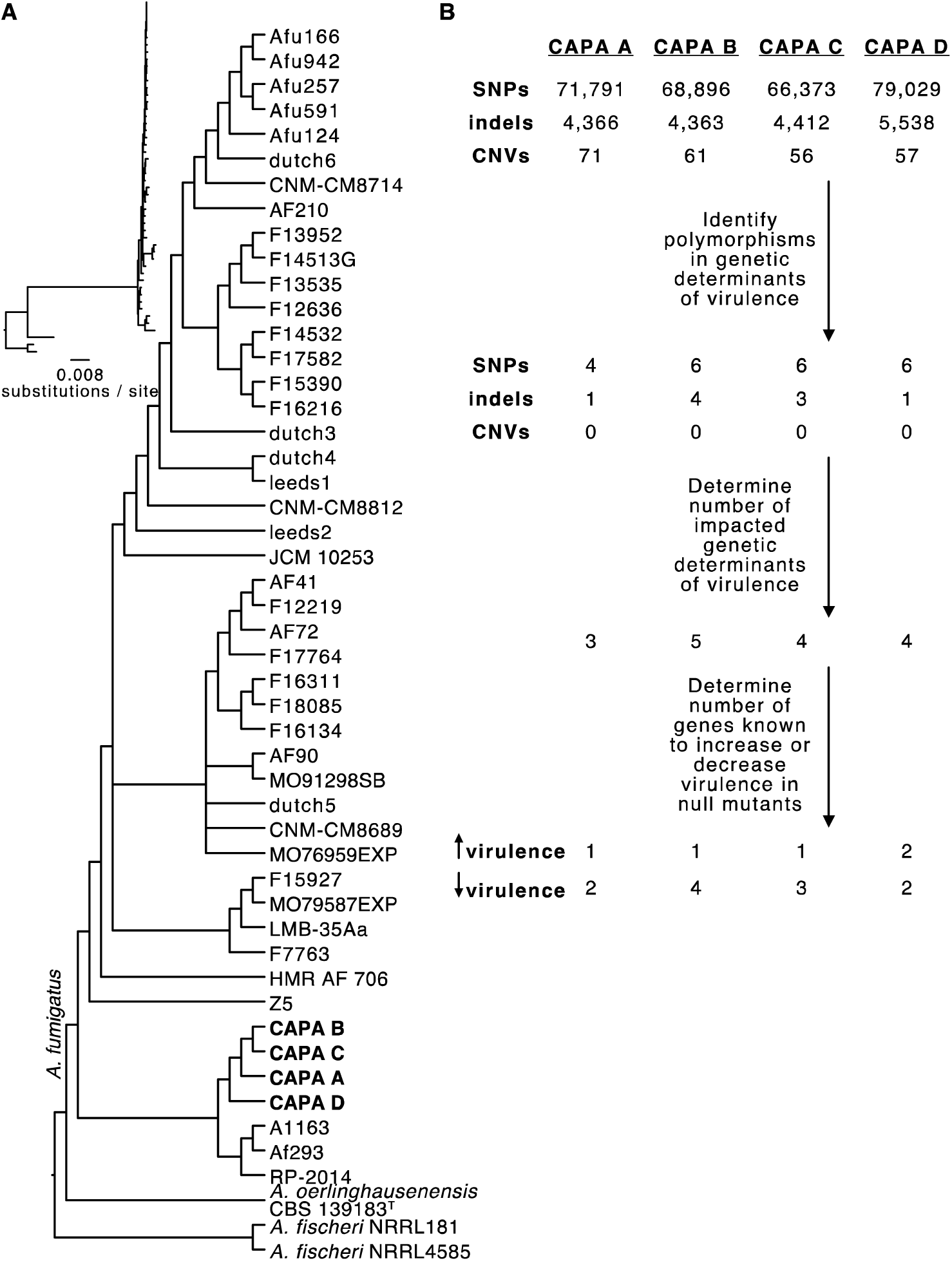
Phylogenomics confirms CAPA isolates are *Aspergillus fumigatus* and mutational spectra among genetic determinants of virulence. (A) Phylogenomic analysis of a concatenated matrix of 4,525 single-copy orthologous groups genes (sites: 7,133,367) confirmed CAPA isolates are *A. fumigatus*. Furthermore, CAPA isolates are closely related to reference strains A1163 and Af293. Bipartitions with less than 85% ultrafast bootstrap approximation support were collapsed. (B) Genome-wide SNPs, indels, and CN variants were filtered for those present in genetic determinants of virulence. Thereafter, the number of genetic determinants of virulence with high impact polymorphisms were identified. The number known to increase or decrease virulence in null mutants was determined thereafter.

### CAPA isolate genomes contain polymorphisms in genetic determinants of virulence and biosynthetic gene clusters

Sequence similarity searches of gene sequences present in a curated list of 206 genetic determinants of virulence previously identified in *A. fumigatus* (File S1) (Abad et al., 2010; Bignell et al., 2016; Kjærbølling et al., 2018; Mead et al., 2019; Urban et al., 2019) showed that all 206 genes were present in the genomes of the CAPA isolates. Furthermore, none of the 206 genetic determinants of virulence showed any copy number variation among CAPA isolates. Examination of single nucleotide polymorphisms (SNPs) and insertion/deletion (indel) polymorphisms coupled with variant effect prediction in these 206 genes (Figure 2B; File S2) showed that all CAPA isolates shared multiple polymorphisms resulting in early stop codons or frameshift mutations suggestive of loss of function (LOF) in *NRPS8* (AFUA_5G12730), a nonribosomal peptide synthetase gene that encodes an unknown secondary metabolite (Lind et al., 2017). LOF mutations in *NRPS8* are known to result in increased virulence in a mouse model of disease (O’Hanlon et al., 2011). Putative LOF mutations were also observed in genes whose null mutants decreased virulence. For example, all CAPA isolates shared the same SNPs resulting in early stop codons that likely result in LOF in *PPTA* (AFUA_2G08590), a putative 4’-phosphopantetheinyl transferase, whose deletion results in reduced virulence in a mouse model of disease (Johns et al., 2017). In light of the close evolutionary relationships among CAPA isolates, we hypothesize that these shared mutations likely occurred in the genome of their most recent common ancestor.

In addition to shared polymorphisms, analyses of CAPA isolate genomes also revealed isolate-specific polymorphisms affecting genetic determinants of virulence (File S2). For example, SNPs resulting in early stop codons, which likely lead to LOF, were observed in *CYP5081A1* (AFUA_4G14780), a putative cytochrome P450 monooxygenase, in CAPA isolates B and C. *CYP5081A1* LOF is associated with reduced virulence of *A. fumigatus* (Mitsuguchi et al., 2009). Additionally, a mutation resulting in the loss of the start codon was observed in *FLEA* (AFUA_5G14740), a gene that encodes an L-fucose-specific lectin, in only CAPA isolate D. Notably, mice infected with *FLEA* null mutants have more severe pneumonia and invasive aspergillosis than wild-type strains. *FLEA* null mutants cause more severe disease because FleA binds to macrophages and therefore is critical for host recognition, clearance, and macrophage killing (Kerr et al., 2016). Most notably, CAPA isolate D had the most putative LOF mutations in the subset of the 206 genetic determinants whose null mutants result in increased virulence, which raises the hypothesis that CAPA isolate D is more virulent than the other CAPA isolates.

Examination of the presence of biosynthetic gene clusters (BGCs) revealed that all CAPA isolates had BGCs that encode secondary metabolites known to modulate host biology (Table 2). For example, all CAPA isolates had BGCs encoding the toxic secondary metabolite gliotoxin (Figure 3). Other intact BGCs in the genomes of the CAPA isolates include fumitremorgin, trypacidin, pseurotin, and fumagillin, which are known to modulate host biology (Ishikawa et al., 2009; González-Lobato et al., 2010; Gauthier et al., 2012); for example, fumagillin is known to inhibit neutrophil function (Fallon et al., 2010, 2011). More broadly, all CAPA isolates had similar numbers and classes of BGCs (Figure S3).

**Table 2.** Biosynthetic gene clusters that produce secondary metabolites implicated in modulating host biology in *A. fumigatus*

|  | Function | Reference(s) | Evidence of Biosynthetic Gene Cluster |  |  |  |
| --- | --- | --- | --- | --- | --- | --- |
|  |  |  | CAPA A | CAPA B | CAPA C | CAPA D |
| Gliotoxin | Inhibits host immune response | (Sugui et al., 2007) | + | + | + | + |
| Fumitremorgin | Inhibits the breast cancer resistance protein | (González-Lobato et al., 2010) | + | + | + | + |
| Trypacidin | Damages lung cell tissues | (Gauthier et al., 2012) | + | + | + | + |
| Pseurotin | Inhibits immunoglobulin E | (Ishikawa et al., 2009) | + | + | + | + |
| Fumagillin | Inhibits neutrophil function | (Fallon et al., 2010, 2011) | + | + | + | + |

‘+’ and ‘-’ indicates the presence and absence of a BGC, respectively.

**Figure 3.**
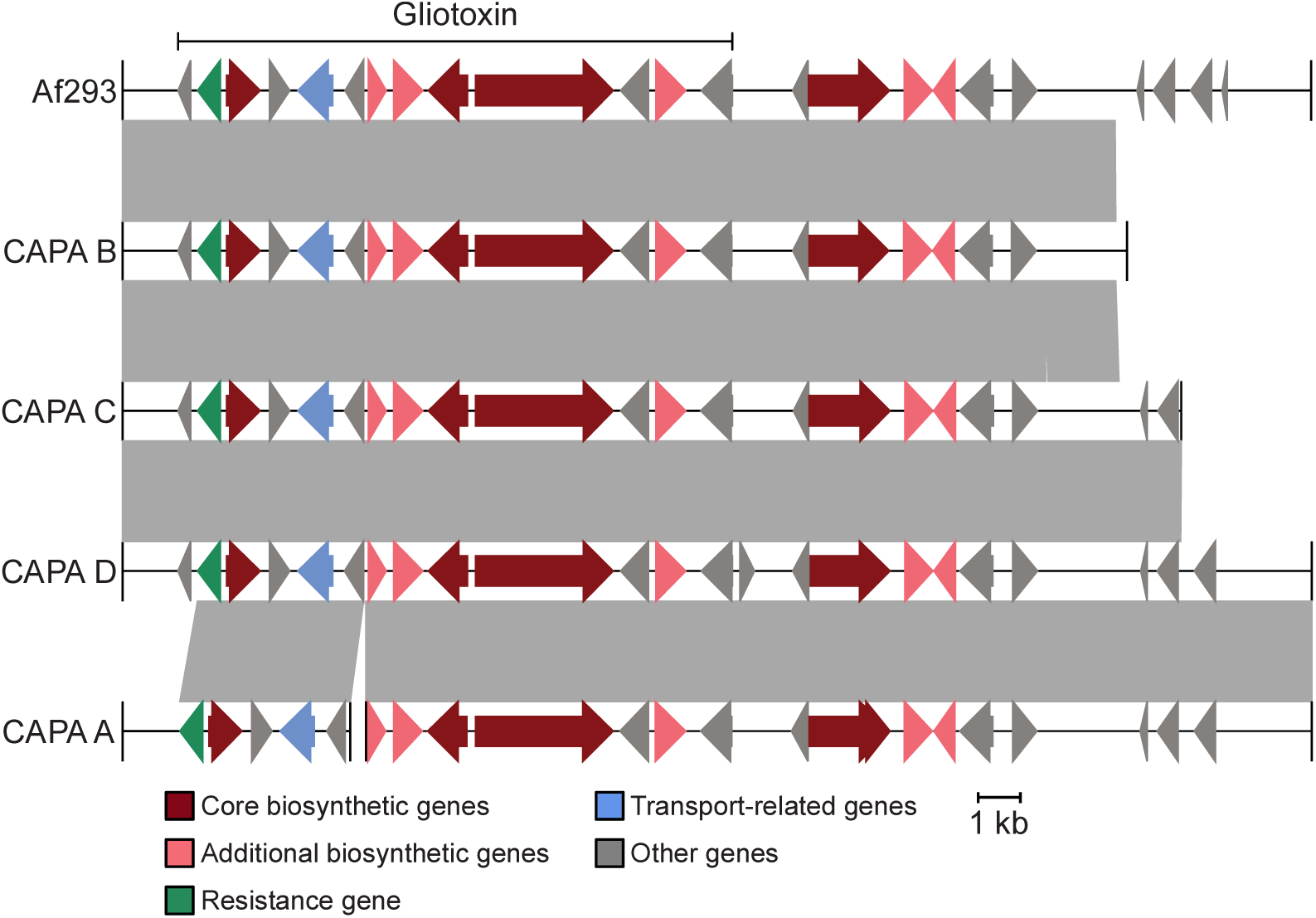
CAPA isolates have BGCs encoding the toxic small molecule gliotoxin. Gliotoxin is known to contribute to virulence of *A. fumigatus*. The genomes of CAPA isolates of *A. fumigatus* contain biosynthetic gene clusters known to encode gliotoxin. Note, the BGC of CAPA A was split between two contigs and therefore the BGC is hypothesized to be present.

In summary, we found that CAPA isolates were closely related to one another and had largely intact genetic determinants of virulence and BGCs. However, we observed strain specific polymorphisms in known genetic determinants of virulence in CAPA isolate genomes, which raises the hypothesis that CAPA isolates differ in their virulence profiles.

### CAPA isolates display strain heterogeneity in virulence but not in virulence-related traits

Examination of virulence and virulence-related traits revealed the CAPA isolates often, but not always, had similar phenotypic profiles compared to reference *A. fumigatus* strains Af293 and CEA17 (which is a *pyrG* mutant derived from the reference strain A1163 (Bertuzzi et al., 2020)). For example, virulence in the *Galleria* moth model of fungal disease revealed strain heterogeneity among CAPA isolates, Af293, and CEA17 (p < 0.001; log-rank test; Figure 4A). Pairwise examination revealed that the observed strain heterogeneity was primarily driven by CAPA isolate D, which was significantly more virulent than all other CAPA isolates (Benjamini-Hochberg adjusted p-value < 0.01 when comparing CAPA isolate D to another isolate; log-rank test; File S3). Also, CAPA isolate C was significantly more virulent than reference strain Af293 (Benjamini-Hochberg adjusted p-value = 0.01; log-rank test). These results reveal that the CAPA isolates have generally similar virulence profiles compared to the reference strains Af293 and CEA17 with the exception of the more virulent CAPA isolate D.

**Figure 4.**
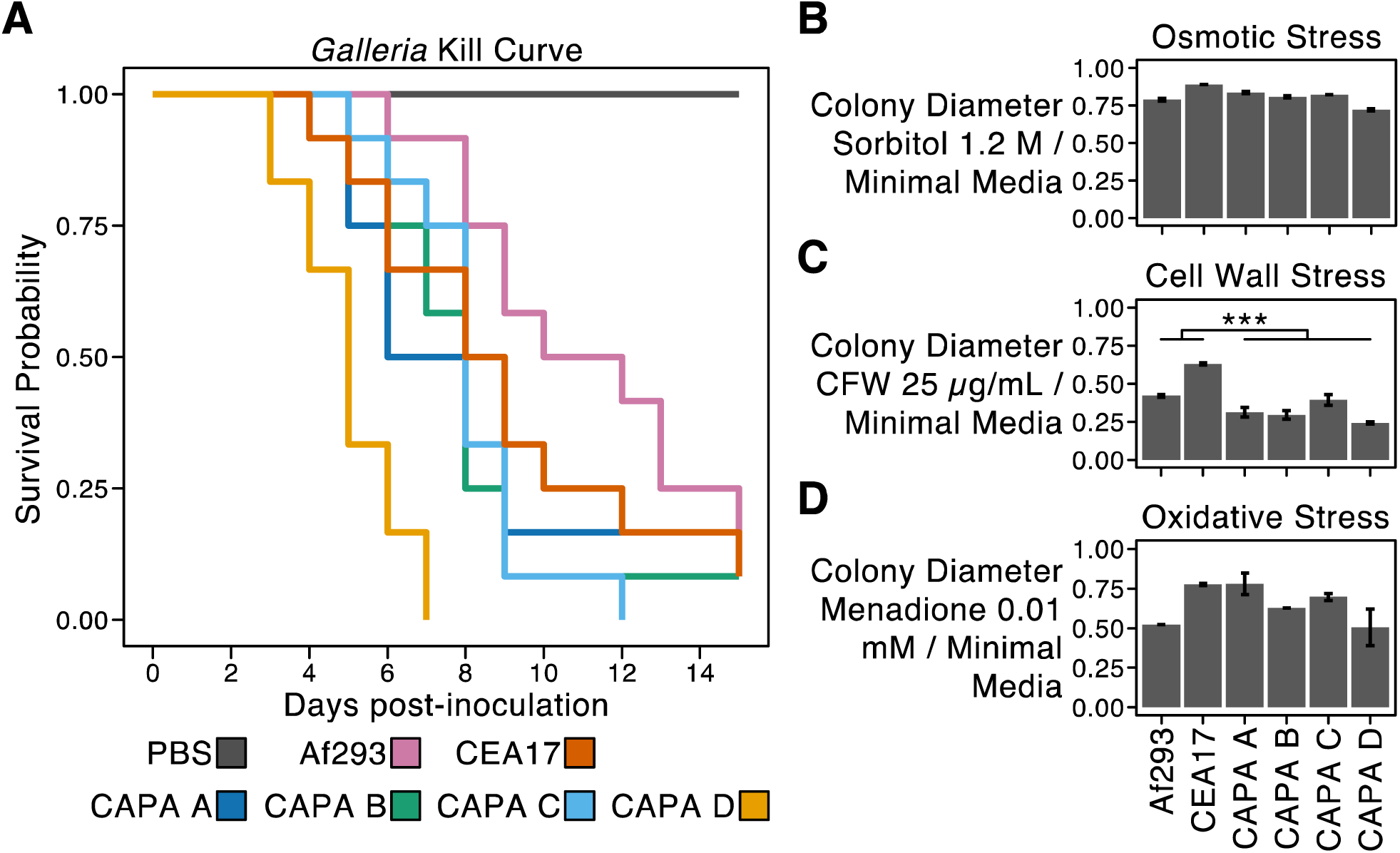
Strain heterogeneity among CAPA isolates. CAPA isolates and reference strains Af293 and CEA17 virulence significantly varied in the *Galleria* moth model of disease (p < 0.001; log-rank test). Pairwise examinations revealed CAPA D was significantly more virulent than all other strains (Benjamini-Hochberg adjusted p-value < 0.01 when comparing CAPA isolate D to another isolate; log-rank test). Growth of CAPA isolates and references strains Af293 and CEA17 in the presence of (B) osmotic, (C) cell wall, and (D) oxidative stressors. Growth differences between CAPA isolates and reference strains Af293 and CEA17 were observed across all growth conditions (p < 0.001; multi-factor ANOVA). Pairwise differences were assessed using the post-hoc Tukey Honest Significant Differences test and were only observed for growth in the presence of CFW at 25 µg/mL (p < 0.001; Tukey Honest Significant Differences test) in which the CAPA isolates did not grow as well as the reference isolates. To correct for strain heterogeneity in growth rates, radial growth in centimeters in the presence of stressors was divided by radial growth in centimeters in the absence of the stressor (MM only). Abbreviations of cell wall stressors are as follows: CFW: calcofluor white; CR: congo red; CSP: caspofungin. Growth in the presence of other stressors is summarized in Supplementary Figure 4.

Examination of growth in the presence of osmotic, cell wall, and oxidative stressors revealed that CAPA isolates had similar phenotypic profiles compared to Af293 and CEA17 (Figures 4B-D and S4) with one exception. Specifically, across all growth assays, we observed significant differences between the CAPA isolates and the reference strains Af293 and CEA17 (p < 0.001; multi-factor ANOVA). Pairwise comparisons revealed significant differences were driven by growth in the presence of calcofluor white wherein the CAPA isolates were more sensitive than reference strains Af293 and CEA17 (p < 0.001; Tukey’s Honest Significant Difference test; Figure 4C). Lastly, antifungal drug susceptibility profiles for amphotericin B, voriconazole, itraconazole, and posaconazole were similar between the CAPA isolates and reference strains Af293 and CEA17 (Table 3).

**Table 3.** Antifungal drug susceptibility of CAPA clinical isolates grown in minimal media

|  | Af293 | CEA17 | CAPA A | CAPA B | CAPA C | CAPA D |
| --- | --- | --- | --- | --- | --- | --- |
| Amphotericin B | 2 | 2 | 2-4 | 2 | 2 | 2 |
| Voriconazole | 1 | 0.25-0.50 | 0.5 | 0.5 | 0.25-0.50 | 0.5 |
| Itraconazole | 0.5 | 0.5 | 0.5 | 0.5 | 0.5 | 0.5 |
| Posaconazole | 1 | 1 | 1 | 1 | 1 | 1 |

Minimum inhibitory concentrations are reported as micrograms (µg) per milliliter (mL).

In summary, we found that the CAPA isolates have similar phenotypic profiles compared to reference strains Af293 and CEA17with the exception of growth in the presence of calcofluor white and the greater virulence of CAPA isolate D. The higher levels of virulence observed in CAPA isolate D may be associated with a greater number of putative LOF mutations that are known to increase virulence; however, this hypothesis requires further functional testing.

### Concluding remarks

The effects of secondary fungal infections in COVID-19 patients are only beginning to be understood. Our results revealed that CAPA isolates are generally, but not always, similar to *A. fumigatus* clinical reference strains. Notably, CAPA isolate D was significantly more virulent than the other three CAPA isolates and two reference strains examined. We hypothesize that this difference is because the D isolate contains more putative LOF mutations in genetic determinants of virulence whose null mutants are known to increase virulence than other strains. Taken together, these results are important to consider in the management of fungal infections among patients with COVID-19, especially those infected with *A. fumigatus*, and broaden our understanding of CAPA.

## Methods

### Patient information and ethics approval

Patients were included into the FungiScope® global registry for emerging invasive fungal infections (www.ClinicalTrials.gov, NCT 01731353). The clinical trial is approved by the Ethics Committee of the University of Cologne, Cologne, Germany (Study ID: 05-102) (Seidel et al., 2017). Since 2019, patients with invasive aspergillosis are also included.

### DNA quality control, library preparation, and sequencing

Sample DNA concentration was measured by Qubit fluorometer and DNA integrity and purity by agarose gel electrophoresis. For each sample, 1-1.5μg genomic DNA was randomly fragmented by Covaris and fragments with average size of 200-400bp were selected by Agencourt AMPure XP-Medium kit. The selected fragments were end-repaired, 3’ adenylated, adapters-ligated, and amplified by PCR. Double-stranded PCR products were recovered by the AxyPrep Mag PCR clean up Kit, and then heat denatured and circularized by using the splint oligo sequence. The single-strand circle DNA (ssCir DNA) products were formatted as the final library and went through further QC procedures. The libraries were sequenced on the MGISEQ2000 platform.

### Genome assembly and annotations

Short-read sequencing data of each sample were assembled using MaSuRCA, v3.4.1 (Zimin et al., 2013). Each *de novo* genome assembly was annotated using the MAKER genome annotation pipeline, v2.31.11 (Holt and Yandell, 2011), which integrates three *ab initio* gene predictors: AUGUSTUS, v3.3.3 (Stanke and Waack, 2003), GeneMark-ES, v4.59 (Besemer and Borodovsky, 2005), and SNAP, v2013-11-29 (Korf, 2004). Fungal protein sequences in the SwissProt database (release 2020_02) were used as homology evidence for the genome annotation. The MAKER annotation process occurs in an iterative manner as described previously (Shen et al., 2018). In brief, for each genome, repeats were first soft-masked using RepeatMasker v4.1.0 (http://www.repeatmasker.org) with the library Repbase library release-20181026 and the “-species” parameter set to “Aspergillus fumigatus”. GeneMark-ES was then trained on the masked genome sequence using the self-training option (“--ES”) and the branch model algorithm (“--fungus”), which is optimal for fungal genome annotation. On the other hand, an initial MAKER analysis was carried out where gene annotations were generated directly from homology evidence, and the resulting gene models were used to train both AUGUSTUS and SNAP. Once trained, the ab initio predictors were used together with homology evidence to conduct a first round of full MAKER analysis. Resulting gene models supported by homology evidence were used to re-train AUGUSTUS and SNAP. A second round of MAKER analysis was conducted using the newly trained AUGUSTUS and SNAP parameters, and once again the resulting gene models with homology supports were used to re-train AUGUSTUS and SNAP. Finally, a third round of MAKER analysis was performed using the new AUGUSTUS and SNAP parameters to generate the final set of annotations for the genome. The completeness of *de novo* genome assemblies and *ab initio* gene predictions was assessed using BUSCO, v4.1.2 (Waterhouse et al., 2018) using 4,191 pre-selected ‘nearly’ universally single-copy orthologous genes from the Eurotiales database (eurotials_odb10.2019-11-20) in OrthoDB, v10.1 (Waterhouse et al., 2013).

### Polymorphism identification

To characterize and examine the putative impact of polymorphisms in the genomes of the CAPA isolates, we identified single nucleotide polymorphisms (SNPs), insertion-deletion polymorphisms (indels), and copy number (CN) polymorphisms. To do so, reads were first quality-trimmed and mapped to the genome of *A. fumigatus* Af293 (RefSeq assembly accession: GCF_000002655.1) following a previously established protocol (Steenwyk and Rokas, 2017). Specifically, reads were first quality-trimmed with Trimmomatic, v0.36 (Bolger et al., 2014), using the parameters leading:10, trailing:10, slidingwindow:4:20, minlen:50. The resulting quality-trimmed reads were mapped to the *A. fumigatus* Af293 genome using the Burrows-Wheeler Aligner (BWA), v0.7.17 (Li, 2013), with the mem parameter. Thereafter, mapped reads were converted to a sorted bam and mpileup format for polymorphism identification using SAMtools, v.1.3.1 (Li et al., 2009).

To identify SNPs and indels, mpileup files were used as input into VarScan, v2.3.9 (Koboldt et al., 2012), with the mpileup2snp and mpileup2indel functions, respectively. To ensure only confident SNPs and indels were identified, a Fischer’s Exact test p-value threshold of 0.05 and minimum variant allele frequency of 0.75 were used. The resulting Variant Call Format files were used as input to snpEff, v.4.3t (Cingolani et al., 2012), which predicted their functional impacts on gene function as high, moderate, or low. To identify CN variants, the sorted bam files were used as input into Control-FREEC, v9.1 (Boeva et al., 2011, 2012). The coefficientOfVariation parameter was set to 0.062 and window size was automatically determined by Control-FREEC. To ensure high-confidence in CN variant identification, a p-value threshold of 0.05 was used for both Wilcoxon Rank Sum and Kolmogorov Smirnov tests.

### Maximum likelihood molecular phylogenetics

To taxonomically identify the species of *Aspergillus* sequenced, we conducted molecular phylogenetic analysis of two different loci and two different datasets. In the first analysis, the nucleotide sequence of the alpha subunit of translation elongation factor EF-1, *tef1* (NCBI Accession: XM_745295.2), from the genome of *Aspergillus fumigatus* Af293 was used to extract other fungal *tef1* sequences from NCBI’s fungal nucleotide reference sequence database (downloaded July 2020) using the blastn function from NCBI’s BLAST+, v2.3.0 (Camacho et al., 2009). *Tef1* sequences were extracted from the CAPA isolates by identifying their best BLAST hit. Sequences from the top 100 best BLAST hits in the fungal nucleotide reference sequence database and the four *tef1* sequences from the CAPA isolates were aligned using MAFFT, v7.402 (Katoh and Standley, 2013) using previously described parameters (Steenwyk et al., 2019) with slight modifications. Specifically, the following parameters were used: --op 1.0 --maxiterate 1000 --retree 1 --genafpair. The resulting alignment was trimmed using ClipKIT, v0.1 (Steenwyk et al., 2020b), with default ‘gappy’ mode. The trimmed alignment was then used to infer the evolutionary history of *tef1* sequences using IQ-TREE2 (Minh et al., 2020). The best fitting substitution model—TIM3 with empirical base frequencies, allowing for a proportion of invariable sites, and a discrete Gamma model (Yang, 1994; Gu et al., 1995) with four rate categories (TIM3+F+I+G4)—was determined using Bayesian Information Criterion. In the second analysis, the same process was used to conduct molecular phylogenetic analysis using calmodulin nucleotide sequences from *Aspergillus* section *Fumigati* species and *Aspergillus clavatus*, an outgroup taxon, using sequences from NCBI that were made available elsewhere (dos Santos et al., 2020a). For calmodulin sequences, the best fitting substitution model was TNe (Tamura and Nei, 1993) with a discrete Gamma model with four rate categories (TNe+G4). Bipartition support was assessed using 5,000 ultrafast bootstrap support approximations (Hoang et al., 2018).

To determine what strains of *A. fumigatus* the CAPA isolates were most similar to, we conducted phylogenomic analyses using the 50 *Aspergillus* proteomes. To do so, we first identified orthologous groups of genes across all 50 *Aspergillus* using OrthoFinder, 2.3.8 (Emms and Kelly, 2019). OrthoFinder takes as input the proteome sequence files from multiple genomes and conducts all-vs-all sequence similarity searches using DIAMOND, v0.9.24.125 (Buchfink et al., 2015). Our input included 50 total proteomes: 47 were *A. fumigatus*, two were *A. fischeri*, and one was *A. oerlinghausenensis* (Fedorova et al., 2008; Lind et al., 2017; Steenwyk et al., 2020d). OrthoFinder then clusters sequences into orthologous groups of genes using the graph-based Markov Clustering Algorithm (van Dongen, 2000). To maximize the number of single-copy orthologous groups of genes found across all input genomes, clustering granularity was explored by running 41 iterations of OrthoFinder that differed in their inflation parameter. Specifically, iterations of OrthoFinder inflation parameters were set to 1.0-5.0 with a step of 0.1. The lowest number ofsingle-copy orthologous groups of genes was 3,399 when using an inflation parameter of 1.0; the highest number was 4,525 when using inflation parameter values of 3.8 and 4.1. We used the groups inferred using an inflation parameter of 3.8.

Next, we built the phylogenomic data matrix and reconstructed evolutionary relationships among the 50 *Aspergillus* genomes. To do so, the protein sequences from 4,525 single-copy orthologous groups of genes were aligned using MAFFT, v7.402 (Katoh and Standley, 2013), with the following parameters: --bl 62 --op 1.0 --maxiterate 1000 --retree 1 --genafpair. Next, nucleotide sequences were threaded onto the protein alignments using function thread_dna in PhyKIT, v0.0.1 (Steenwyk et al., 2020a). The resulting codon-based alignments were then trimmed using ClipKIT, v0.1 (Steenwyk et al., 2020b), using the gappy mode. The resulting aligned and trimmed alignments were then concatenated into a single matrix with 7,133,367 sites using the PhyKIT function create_concat. To reconstruct the evolutionary history of the 50 *Aspergillus* genomes, a single best-fitting model of sequence substitution and rate heterogeneity was estimated across the entire matrix using IQ-TREE2, v.2.0.6 (Minh et al., 2020). The best-fitting model was determined to be a general time reversible model with empirical base frequencies and invariable sites with a discrete Gamma model with four rate categories (GTR+F+I+G4) (Tavaré, 1986; Gu et al., 1995; Waddell and Steel, 1997; Vinet and Zhedanov, 2011) using Bayesian Information Criterion. During tree search, the number of candidate trees maintained during maximum likelihood tree search was increased from five to ten. Five independent searches were conducted and the tree with the best log-likelihood score was chosen as the ‘best’ phylogeny. Bipartition support was evaluated using 5,000 ultrafast bootstrap approximations (Hoang et al., 2018).

### Biosynthetic gene cluster prediction

To predict BGCs in the genomes of *A. fumigatus* strains Af293 and the CAPA isolates, gene boundaries inferred by MAKER were used as input into antiSMASH, v4.1.0 (Weber et al., 2015). Using a previously published list of genes known to encode BGCs in the genome of *A. fumigatus* Af293 (Lind et al., 2017), BLAST-based searches using an expectation value threshold of 1e-10 were used to identify BGCs implicated in modulating host biology using NCBI’s BLAST+, v2.3.0 (Camacho et al., 2009). Among predicted BGCs that did not match the previously published list, we further examined their evolutionary history if at least 50% of genes showed similarity to species outside of the genus *Aspergillus*, which is information provided in the antiSMASH output. Using these criteria, no evidence suggestive of horizontally acquired BGCs from distant relatives was detected.

### Infection of *Galleria mellonella*

Survival curves (n=12/strain) of *Galleria mellonella* infected with CAPA isolates A, B, C, and D. Phosphate buffered saline (PBS) without asexual spores (conidia) was administered as a negative control. A log-rank test was used to examine strain heterogeneity followed by pairwise comparisons with Benjamini-Hochberg multi-test correction (Benjamini and Hochberg, 1995). All the selected larvae of *Galleria mellonella* were in the final (sixth) instar larval stage of development, weighing 275–330 milligram. Fresh conidia from each strain were harvested from minimal media (MM) plates in PBS solution and filtered through a Miracloth (Calbiochem). For each strain, the spores were counted using a hemocytometer and the stock suspension was done at 2 × 10^8^ conidia/milliliter. The viability of the administered inoculum was determined by plating a serial dilution of the conidia on MM medium at 37°C. A total of 5 microliters (1 × 10^6^ conidia/larva) from each stock suspension was inoculated per larva. The control group was composed of larvae inoculated with 5 microliters of PBS to observe the killing due to physical trauma. The inoculum was performed by using Hamilton syringe (7000.5KH) via the last left proleg. After infection, the larvae were maintained in petri dishes at 37°C in the dark and were scored daily. Larvae were considered dead by presenting the absence of movement in response to touch.

### Growth assays

To examine growth conditions of the CAPA isolates and reference strains Af293 and A1163, plates were inoculated with 10^4^ spores per strain and allowed to grow for five days on solid MM or MM supplemented with various concentrations of osmotic (sorbitol, NaCl), cell wall (congo red, calcofluor white and caspofungin), and oxidative stress agents (menadione and t-butyl) at 37°C. MM had 1% (weight / volume) glucose, original high nitrate salts, trace elements, and a pH of 6.5; trace elements, vitamins, and nitrate salts compositions follow standards described elsewhere (Käfer, 1977). To correct for strain heterogeneity in growth rates, radial growth in centimeters in the presence of stressors was divided by radial growth in centimeters in the absence of the stressor.

### Data Availability

Newly sequenced genomes assemblies, annotations, and raw short reads have been deposited to NCBI’s GenBank database under BioProject accession PRJNA673120. Additional copies of genome assemblies, annotations, and gene coordinates have been uploaded to figshare (doi: 10.6084/m9.figshare.13118549). Other raw data including the genome assembly and annotations of all analyzed *Aspergillus* genomes, the aligned and trimmed phylogenetic and phylogenomic data matrices, predicted BGCs, and other analysis have been uploaded to figshare as well.

## Conflict of Interest

Oliver A. Cornely is supported by the German Federal Ministry of Research and Education, is funded by the Deutsche Forschungsgemeinschaft (DFG, German Research Foundation) under Germany’s Excellence Strategy – CECAD, EXC 2030 – 390661388 and has received research grants from, is an advisor to, or received lecture honoraria from Actelion, Allecra Therapeutics, Al-Jazeera Pharmaceuticals, Amplyx, Astellas, Basilea, Biosys, Cidara, Da Volterra, Entasis, F2G, Gilead, Grupo Biotoscana, IQVIA, Janssen, Matinas, Medicines Company, MedPace, Melinta Therapeutics, Menarini, Merck/MSD, Mylan, Nabriva, Noxxon, Octapharma, Paratek, Pfizer, PSI, Roche Diagnostics, Scynexis, and Shionogi. Philipp Koehler has received non-financial scientific grants from Miltenyi Biotec GmbH, Bergisch Gladbach, Germany, and the Cologne Excellence Cluster on Cellular Stress Responses in Aging-Associated Diseases, University of Cologne, Cologne, Germany, and received lecture honoraria from or is advisor to Akademie für Infektionsmedizin e.V., Astellas Pharma, European Confederation of Medical Mycology, Gilead Sciences, GPR Academy Ruesselsheim, MSD Sharp & Dohme GmbH, Noxxon N.V., and University Hospital, LMU Munich outside the submitted work. Antonis Rokas is a Scientific Consultant for LifeMine Therapeutics, Inc.

## Acknowledgements

We thank the Rokas and Goldman laboratories for support of this work and their helpful insight. J.L.S. and A.R. are supported by the Howard Hughes Medical Institute through the James H. Gilliam Fellowships for Advanced Study Program. A.R. received additional support from a Discovery grant from Vanderbilt University, the Burroughs Wellcome Fund, the National Science Foundation (DEB-1442113), and National Institutes of Health/National Institute of Allergy and Infectious Diseases (1R56AI146096-01A1). G.H.G. and A.D. thank Fundação de Amparo à Pesquisa do Estado de São Paulo (FAPESP) grant numbers 2016/07870-9 and 2020/06151-4, and Conselho Nacional de Desenvolvimento Científico e Tecnológico (CNPq), both from Brazil. X.Z. is supported by the Key-Area Research and Development Program of Guangdong Province (2018B020206001). F.F. has a Clinician Scientist Position supported by the Deans Office, Faculty of Medicine, University of Cologne. C.V. is supported by FAPESP grant number 2018/00715-3. S.L.K. was supported by the National Center for Complementary and Integrative Health, a component of the National Institutes of Health, under award number F31 AT010558.

## Notes

https://doi.org/10.6084/m9.figshare.13118549.v1

