## Supplementary Figures for "Genomic and phenotypic analysis of COVID-19-associated pulmonary aspergillosis isolates of *Aspergillus fumigatus*"

Table S1. Genome assembly quality and statistics

|  | CAPA A | CAPA B | CAPA C | CAPA D |
| --- | --- | --- | --- | --- |
| Genome Size | 29,072,951 | 28,651,837 | 28,214,156 | 28,390,836 |
| Number of Genes | 9,919 | 9,697 | 9,686 | 9,672 |
| Number of Scaffolds | 356 | 381 | 295 | 466 |
| N50 | 493,663 | 467,188 | 459,253 | 315,890 |
| Depth of Coverage | 176.6 | 193.8 | 200.3 | 180.5 |
| BUSCO stats | 4161, 15, 2, 13;  99.28, 0.36, 0.48, 0.31 | 4167, 10, 1, 13;  99.43, 0.24, 0.02, 0.31 | 4165, 4, 3, 19;  99.38, 0.10, 0.07, 0.45 | 4168, 7, 2, 21;  99.45, 0.17, 0.5, 0.50 |

BUSCO stats are reported using the following convention: single copy, duplicated, fragmented, and missing genes, in which the first row is the number of genes and the second row is the percentages.

**Supplementary figure 1. Molecular phylogenetics of fungal *tef1* sequences reveals CAPA isolates are *Aspergillus fumigatus*.** (A) Using the *tef1* sequences from the genome of *A. fumigatus*, the top 100 best BLAST hits of *tef1* sequences from the fungal reference sequence database were combined with the *tef1* sequences from the CAPA isolates. After multiple sequence alignment and trimming, their evolutionary history was inferred using maximum likelihood phylogenetics. (B) The blue circle indicates were *Aspergillus* sequences are found on the tree and the CAPA isolates are closely related to *A. fumigatus* (strain Af293). Red lines indicate bipartitions with full support. (C) To provide a clearer view of the relationship of *Aspergillus* sequences, a cladogram reveals the CAPA isolates further highlight that the CAPA isolates are closely related to *A. fumigatus*. Bipartition support values greater than 95 are shown.

**Supplementary figure 2. Molecular phylogenetics of calmodulin sequences from section *Fumigati* species confirms CAPA isolates are *Aspergillus fumigatus*.** Calmodulin sequences from species from section *Fumigati* were acquired from NCBI, supplemented with calmoduin sequences from the CAPA isolates, aligned, trimmed, and their evolutionary history was inferred using maximum likelihood phylogenetics. Examination of the resulting phylogeny revealed the CAPA isolates (in bold) are nested with a clade of other *A. fumigatus* isolates. The ancestral bipartition of this lineage received full bipartition support. Bipartition support was inferred using ultrafast bootstrap support as implemented in IQ-TREE2. Bipartition support greater than 90 is shown on the phylogeny. Upper left inset uses meaningful branch lengths while the large phylogeny is a cladogram.

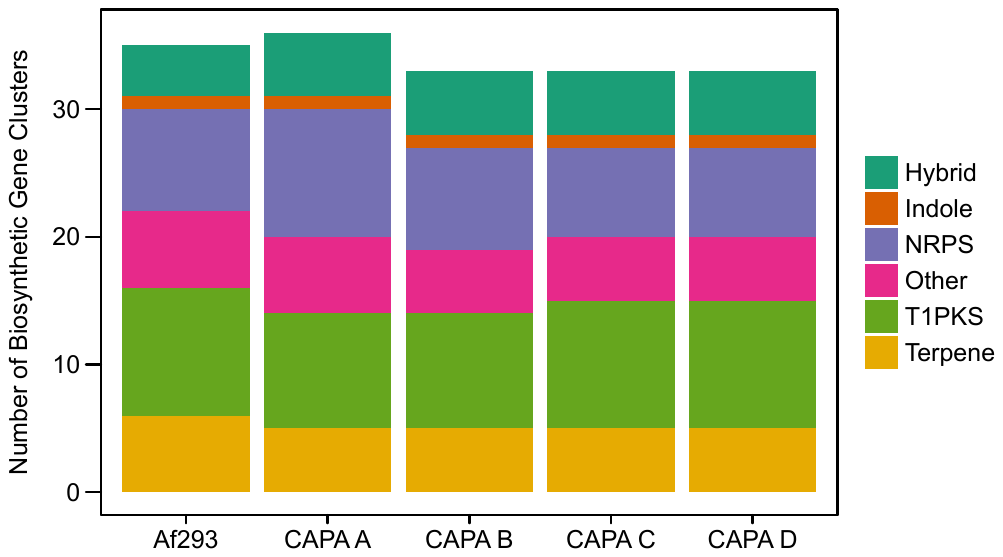

**Supplementary figure 3. CAPA isolates of *Aspergillus fumigatus* have similar numbers and types of biosynthetic gene clusters as Af293.** Examination of the number and types of biosynthetic gene clusters in CAPA isolates of *Aspergillus fumigatus* reveal they have a similar types and numbers of BGCs. Classes of BGCs are written using the following

abbreviations: NRPS: nonribosomal peptide synthetase; T1PKS: type I polyketide synthase;

Hybrid: a combination of multiple BGC classes.

**Supplementary figure 4. Growth of CAPA isolates in cell wall, oxidative, and osmotic stressor do not differ from reference isolates Af293 and CEA17.** Growth of CAPA isolates and references strains Af293 and CEA17 in the presence of (A) cell wall, (B) oxidative, and (C) osmotic stressors. Growth differences between CAPA isolates and references isolates Af293 and CEA17 were observed across all growth conditions (p < 0.001; multi-factor ANOVA). Pairwise differences were assessed using the post-hoc Tukey Honest Significant Differences (THSD) test. Significant differences were observed for growth in the presence of the cell wall stressor CFW (p < 0.001; THSD test; see main figure 4) and no other condition. Abbreviations of cell wall stressors are as follows: CFW: calcofluor white; CR: congo red; CSP: caspofungin. To correct for strain heterogeneity in growth rates, radial growth in centimeters in the presence of stressors was divided by radial growth in centimeters in the absence of the stressor (MM only).
